# Nitrogen- and phosphorus-starved *Triticum aestivum* show distinct belowground microbiome profiles

**DOI:** 10.1101/507863

**Authors:** Antoine P. Pagé, Julien Tremblay, Luke Masson, Charles W. Greer

**Affiliations:** Aquatic and Crop Resource Development Research Centre, National Research Council Canada, Montréal, QC, Canada.; Energy, Mining and Environment Research Centre, National Research Council Canada, Montréal, QC, Canada.; Human Health Therapeutics Research Centre, National Research Council Canada, Montréal, QC, Canada.

## Abstract

Many plants have natural partnerships with microbes that can boost their nitrogen (N) and/or phosphorus (P) acquisition. To assess whether wheat may have undiscovered associations of these types, we tested if N/P-starved *Triticum aestivum* show microbiome profiles that are simultaneously different from those of N/P-amended plants and those of their own bulk soils. The bacterial and fungal communities of root, rhizosphere, and bulk soil samples from the Historical Dryland Plots (Lethbridge, Canada), which hold *T. aestivum* that is grown both under N/P fertilization and in conditions of extreme N/P-starvation, were taxonomically described and compared (bacterial 16S rRNA genes and fungal Internal Transcribed Spacers - ITS). As the list may include novel N- and/or P-providing wheat partners, we then identified all the operational taxonomic units (OTUs) that were proportionally enriched in one or more of the nutrient starvation- and plant-specific communities. These analyses revealed: a) distinct N-starvation root and rhizosphere bacterial communities that were proportionally enriched, among others, in OTUs belonging to families *Enterobacteriaceae, Chitinophagaceae, Comamonadaceae, Caulobacteraceae, Cytophagaceae, Streptomycetaceae*, b) distinct N-starvation root fungal communities that were proportionally enriched in OTUs belonging to taxa *Lulworthia, Sordariomycetes, Apodus, Conocybe, Ascomycota, Crocicreas*, c) a distinct P-starvation rhizosphere bacterial community that was proportionally enriched in an OTU belonging to genus *Agrobacterium*, and d) a distinct P-starvation root fungal community that was proportionally enriched in OTUs belonging to genera *Parastagonospora* and *Phaeosphaeriopsis*. Our study might have exposed wheat-microbe connections that can form the basis of novel complementary yield-boosting tools.

## Introduction

The spread of chemical fertilization practices was one of the main features of the Green Revolution. These yield-boosting methods are so efficient that they quickly became worldwide pillars of intensive agriculture. The environmental costs associated with the currently generalized use of inorganic N and P fertilizers are, however, mounting [1–4]. Although it remains manageable now, this imbalance could eventually compromise the sustainability of our farming system [5–6]. It therefore constitutes a strong incentive for the development of complementary yield-boosting tools.

Farmers already have access to many such tools, including crop cultivars that possess improved nutrient use efficiencies, alternative agronomic practices, and microbial inoculants [7–10]. However, although appreciable, the reduction of global environmental impacts that these tools can collectively provide appears limited [9, 11]. More will likely be needed to address the future challenges of sustainable food production. As many of the currently deployed complementary yield-boosting tools rest on natural N– or P-sharing plant-microbe partnerships, discovering new partnerships of these types could help us develop the necessary tools.

Targeting wheat may be a good choice for research endeavors that have such objectives. Several studies suggest that the plant can partner with N- and P-providing microorganisms [12–20], but its natural microbial associations are not well delineated. Since wheat farming currently consumes approximately 20% of the worldwide production of inorganic N and P fertilizers [21], this choice also opens up the possibility of producing highly valuable complementary yield-boosting tools. We thus focused the work presented herein on *T. aestivum*, the most widely grown wheat species.

Correlations between microbiome composition and soil N/P content have been reported for many plants [22–32]. This phenomenon is largely attributed to concomitant variations of soil microbiome composition, which presumably prompt different unsolicited plant colonization processes [33, 34]. But the dynamics of N- and P-sharing plant-microbe partnerships are undoubtedly also at play. The N-related microbiome variations seen in legumes are, for example, largely tied to the plants’ modulation of their rhizobia recruitment efforts [35–37]. The P-related microbiome variations seen in many mycorrhizal embroyphytes are similarly tied to the recruitment of arbuscular mycorrhizae [38–40]. The prevalence of N/P-sharing plant-microbe partnerships, and their contribution to N/P-related variations of plant microbiome composition, seem to extend much beyond the above examples [41–45]. To assess whether wheat may be among the plants that have undiscovered associations of these types, we thus first aimed to test if its root and rhizosphere microbiomes vary with soil N and P content.

The microbiomes of plant roots and rhizospheres are generally distinguishable from those of the adjacent bulk soils [46]. This phenomenon is again largely attributed to unsolicited microbial colonization. But plant recruitment of N- and P-providing microorganisms is also undoubtedly at play under conditions of low soil N and P. Legumes’ activation of flavonoid release and nodule organogenesis mechanisms, for example, promotes such contrasts [36]. Similar patterns are also attributed to other microbial attraction and accommodation mechanisms [42, 44]. We thus, as part of the aforementioned assessment, also aimed to test if wheat shapes its root and rhizosphere microbiomes under conditions of low N and P availability.

The Historical Dryland Plots, a unique long-term experiment that examines wheat production with and without the use of N and P fertilizers in Lethbridge, Canada [47], constituted the ideal setting to run the selected tests. Indeed, its Rotation A section (wheat without fallow since 1911) contains a plot that annually receives N and P fertilization (45 kg N ha^-1^ ammonium nitrate and 20 kg P ha^-1^ monocalcium phosphate, i.e. N_45_P_20_) and plots that have not been amended with one or both for over 4 decades (i.e. N_45_P_0_, N_0_P_20_, N_0_P_0_). It thus offers an access to *T. aestivum* that are both grown under sufficient N/P supplies and in conditions of extreme N/P-starvation on the same land. We therefore profiled the bacterial and fungal communities that were associated with *T. aestivum* roots, rhizospheres, and bulk soil collected from each Rotation A plot, and tested: 1) if the nutrient-starved *T. aestivum* showed root and rhizosphere microbial communities that were significantly different from those of the fertilized plants (hypothesis 1, Fig 1), and 2) if starvation-specific *T. aestivum* communities were also significantly different from those of the adjacent bulk soils (hypothesis 2, Fig 1). As they may represent soil microbes that are specifically attracted towards the plant when it experiences N– or P-starvation, a behavior that is seen in known N- and P-providing plant partners [46, 48, 49], we then identified all the OTUs that were proportionally enriched in one or more of the nutrient starvation- and plant-specific communities.

**Fig 1.**
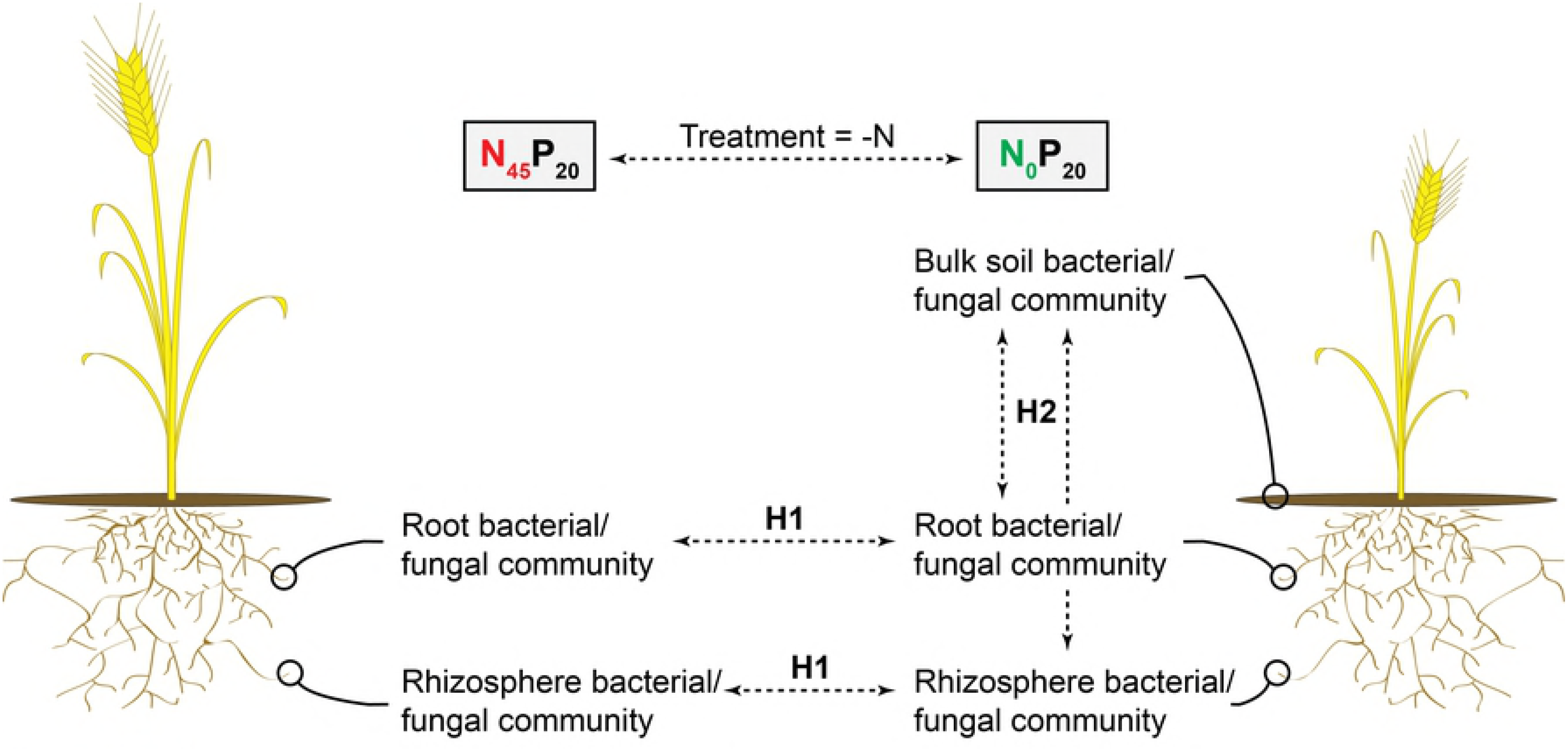
Microbiome comparison procedure. Procedure used herein to identify starvation- and *T. aestivum*-specific bacterial and fungal communities (example illustrating the assessment of N-starved *T. aestivum* communities). On the left, a *T. aestivum* plant grown with N fertilization. On the right, a *T. aestivum* plant grown without N fertilization. H1 = hypothesis 1: the nutrient-starved *T. aestivum* show root and rhizosphere microbial communities that are significantly different from those of the fertilized plants. H2 = hypothesis 2: starvation-specific *T. aestivum* communities are significantly different from those of the adjacent bulk soil.

**Fig 2.**
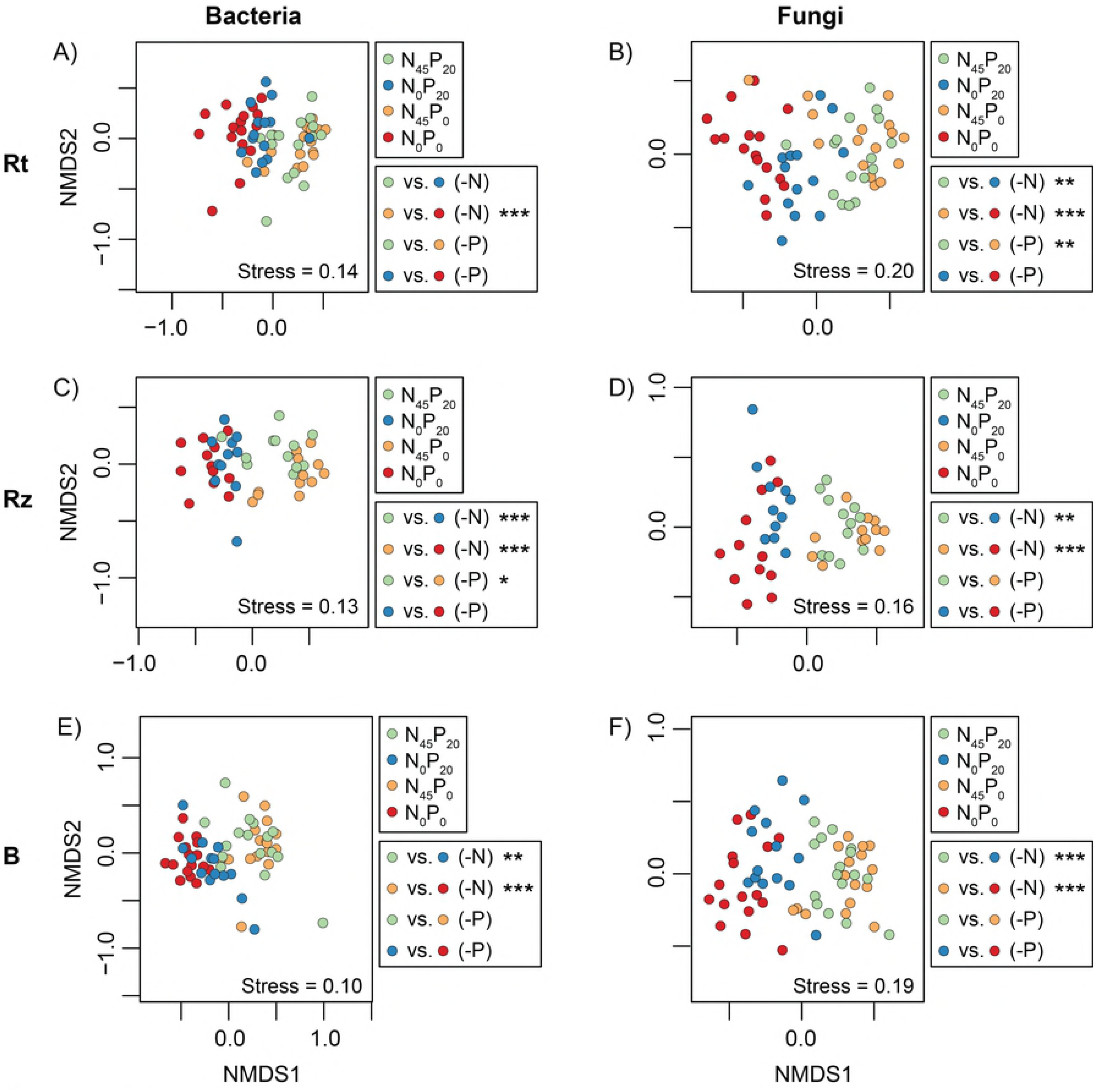
Microbial community comparisons - fertilization. Non-metric multidimensional scaling (NMDS) ordinations representing the dissimilarity (Bray-Curtis) of microbial communities sampled in the different Rotation A plots. The significance of pairwise comparisons is reported next to each graph (* = p < 0.05; ** = p < 0.01, *** = p < 0.001).

## Material and methods

### Sampling and DNA extraction

DNA samples analyzed in this study were provided by B. Helgason (Agriculture and Agri-Food Canada and University of Saskatchewan) and S. Hemmingsen (National Research Council Canada). Plant and soil samples (S1 Table) were collected from the Rotation A plot of the Rotation ABC study at the AAFC Lethbridge Research Center in 2013 and 2014, when the plants were in early vegetative growth and anthesis, following the procedure described by Siciliano and Germida [50]. DNA was extracted using DNeasy PowerSoil and PowerPlant Pro DNA Isolation Kits (Qiagen, Hilden, Germany) from bulk soil (250 mg per sample), rhizosphere soil (250 mg per sample), and washed root samples (50 mg per sample) collected by Helgason (Hemmingsen personal communication). Rotation A plots N_45_P_0_, N_0_P_20_, and N_45_P_20_ were supplemented once a year with 45 kg N ha-1 ammonium nitrate and/or 20 kg P ha-1 monocalcium phosphate [47]. AC Lillian was planted during the period under study.

### PCR amplification/sequencing of microbial taxonomic markers

Genes encoding for a portion of the V4 region of bacterial 16S rRNA were amplified from DNA extracts by PCR, with the following reagents and conditions: 1X HotStarTaq Plus Master Mix (Qiagen, Hilden, Germany), 0.4 mg/mL bovine serum albumin (Roche Diagnostics, Basel, Switzerland), and 0.6 uM each of forward (F520 5’-TCG TCG GCA GCG TCA GAT GTG TAT AAG AGA CAG AGC AGC CGC GGT AAT-3’) and reverse (R799 5’-GTC TCG TGG GCT CGG AGA TGT GTA TAA GAG ACA GCA GGG TAT CTA ATC CTG TT-3’, based on Huws et al. [51] MiSeq primers (Integrated DNA Technologies, Coralville, USA) per reaction; 5 min at 95°C, followed by 25 cycles of 30 sec at 95°C, 30 sec annealing at 45°C, 45 sec at 72°C, and a final 10 min at 72°C. Fungal internal transcribed spacers were also PCR amplified using the same reagents and conditions as bacterial 16S rRNA genes, except for the forward (ITS_F 5’-TCG TCG GCA GCG TCA GAT GTG TAT AAG AGA CAG CTT GGT CAT TTA GAG GAA GTA A -3’) and reverse (ITS_R 5’-GTC TCG TGG GCT CGG AGA TGT GTA TAA GAG ACA GCT GCG TTC TTC ATC GAT -3’, based on Martin and Rygiewicz [52]) primers.

PCR amplicons were purified with Beckman Coulter’s Agencourt AMPure XP system (Beckman Coulter Inc, Brea, USA) according to the manufacturer’s instructions. They were then indexed by PCR with the following reagents and conditions: 5 uL of purified amplicon solution, 1X KAPA HiFi HotSart ReadyMix (Kapa Biosystems, Wilmington, USA), and 2.5 uL each of Nextera XT DNA Library Prep (Illumina, San Diego, USA) index 1 and index 2 primer; 3 min at 95°C, followed by 8 cycles of 30 sec at 95°C, 30 sec annealing at 55°C, 30 sec at 72°C, and a final 5 min at 72°C. Indexed PCR amplicons were again purified using Beckman Coulter’s Agencourt AMPure XP system, quantified with a Quant-iT PicoGreen dsDNA Assay Kit (ThermoFisher Scientific, Waltham, USA), pooled, size selected with a SPRISelect DNA Size Selection system (Beckman Coulter Inc, Brea, USA), and sequenced on a MiSeq DNA Sequencer (Illumina, San Diego, USA). Sequence data was deposited into the NCBI’s sequence read archive under bioproject PRJNA343655 and biosamples SAMN05791727 to SAMN05791930.

### Sequence quality control, collation of sequences into OTUs, normalization of OTU abundances, and assignment of taxonomic qualifiers

Sets of 16S rRNA gene and ITS sequence reads were all analyzed with a custom amplicon analysis pipeline [53]. Briefly: common sequence contaminants were removed from raw reads using a kmer matching tool (DUK v1.051, http://duk.sourceforge.net/), filtered reads were assembled using FLASH v1.2.11 [54], assembled amplicons were trimmed with custom Perl scripts to remove remaining primer sequences and filtered for quality (sequencing with >1 N, an average quality score lower than 33, or more than 5 nucleotides having a quality score lower than 10 were rejected).

OTU generation was conducted using a three-step clustering pipeline. Quality controlled sequences were dereplicated at 100% identity using a custom Perl script, denoised at 99% identity using USEARCH v.6.0.203 [55]. Clusters of less than three reads were discarded and remaining clusters were scanned for chimeras using UCHIME, first in *de novo* mode, then in reference mode, using the Broad Institute’s 16S rRNA Gold reference database. Remaining clusters were clustered at 97% identity (USEARCH).

16S rRNA gene OTUs were assigned taxonomic qualifiers using the RDP classifier (v2.5) with a modified Greengenes training set built from a concatenation of the Greengenes database (version 13_8 maintained by Second Genome), Silva eukaryotes 18S r118, and a selection of chloroplast and mitochondrial rRNA sequences. For ITS OTUs, this task was performed using the ITS Unite database. Hierarchical tree files were generated with custom Perl scripts and used to generate training sets using the RDP classifier training set generator’s functionality [56]. With taxonomic qualifiers in hand, OTU abundances were normalized with edgeR v3.10.2 [57].

### Estimated microbial community coverages, comparisons of community structures, identification of OTU enrichments

The proportion of each microbial community that was captured through the sequencing efforts described above was estimated using R package Entropart (Chao estimator) [58].

We then sought to determine whether significant differences of mean community dissimilarity were found among the bacterial root data of the four Rotation A plots (N_0_P_0_, N_0_P_20_, N_45_P_0_, N_45_P_20_). This was accomplished by running a permanova analysis on a matrix of community dissimilarity (Bray-Curtis index) created with the associated sample profiles (9999 permutations, R package Vegan version 2.4-1) [59]. This procedure was subsequently repeated with bacterial rhizosphere, bacterial bulk soil, fungal root, fungal rhizosphere, and fungal bulk soil data. Datasets that showed significant differences (p < 0.05) were further explored with post hoc permanova analyses that compared plot-specific data subsets against each other (e.g. bacterial root N_45_P_20_ vs. N_0_P_20_ and N_45_P_0_ vs. N_0_P_0_ for N starvation, bacterial root N_45_P_20_ vs. N_0_P_0_ and N_0_P_20_ vs. N_0_P_0_ for P starvation, hypothesis 1 - Fig 1, analyses parameters as above). The p values of these individual pairwise comparisons were adjusted with the Bonferroni correction [60] to account for multiple testing. The tests that revealed significant differences (p < 0.05 after adjustment) were followed by tests for multivariate homogeneity of group dispersion conducted with Vegan function betadisper and R stats package function tukeyHSD (default parameters, p values also adjusted with Bonferroni correction). Non-metric multidimensional scaling (NMDS) ordinations were performed with Vegan’s metaMDS function to illustrate community differences (trymax = 100, all other parameters default).

We then sought to determine whether the identified starvation-specific communities were also significantly different from their bulk soil counterparts (e.g. bacterial root N_0_P_0_ vs. bacterial bulk soil N_0_P_0_, hypothesis 2 - Fig 1). This was also accomplished by running permanova analyses on matrices of community dissimilarity (parameters as above with Bonferroni corrections). The tests that revealed significant differences (p < 0.05 after adjustment) were again followed by tests for multivariate homogeneity of group dispersion (parameters as above), and community differences were again illustrated with NMDS ordinations.

These analyses identified 6 N/P-starvation and *T. aestivum*-specific microbial communities (i.e. that were simultaneously different from the communities of fertilized *T. aestivum* and those of their adjacent bulk soil). A conservative two-step process was used to identify the OTUs that were enriched in each one. All OTUs that were detected in one or more of the considered compartment- and plot-specific data subsets (e.g. bacterial root N_0_P_0_, bacterial root N_45_P_0_, bacterial bulk soil N_0_P_0_,) were, in each case, individually submitted to a two-tailed Kruskal-Wallis Rank Sum Test using function kruskal.test from R stats package (default parameters). Those that had a significantly different average relative abundance among the three data subsets were then assessed for statistical enrichments in the starvation- and *T. aestivum*-specific data subset (e.g. bacterial root N_0_P_0_). This was accomplished by fitting a linear model to each OTU’s overall abundance in the three considered data subsets (using R stats package lm function), and repeatedly testing it for abundance differences in pairs of compartment- and plot-specific data subsets (e.g. bacterial root N_0_P_0_ vs bacterial root N_45_P_0_, bacterial root N_0_P_0_ vs bacterial bulk soil N_0_P_0_) using R package Multcomp’s general linear hypotheses testing function (glht) with the treatment difference set to “Tukey” [61]. To illustrate the results of these analyses, we presented the average relative abundances of each enriched OTU next to a dendrogram built with the average Bray-Curtis dissimilarities of the associated compartment- and plot-specific data subsets. Heatmap were drawn using R package pheatmap [62], average community similarities were calculated using Vegan’s meandist function, and dendrograms were drawn with function of hclust of R’s stats package.

## Results

### Estimated microbial community coverages, comparisons of community structures, identification of OTU enrichments

The sequencing datasets associated with the 172 samples contained, on average, 50052 bacterial 16S rRNA gene sequences or 77115 fungal ITS sequences following quality control. Evaluations of community coverage calculated using a Chao estimator suggest that this sequencing effort allowed the identification of, on average, 98.1% of the bacterial OTUs and 99.4% of the fungal OTUs that were present in each sample.

Significant differences of mean community dissimilarity were found among the bacterial root data of the four Rotation A plots (p = 0.0001, R^2^ = 0.13), and the difference found in a pair of N-starved/N-fertilized data subsets contributed to this signal (N_45_P_0_/N_0_P_0_, p = 0.0004, R^2^ = 0.13, Fig 2A). An additional comparison revealed that the N_0_P_0_ root data subset was also significantly different from that of N_0_P_0_’s bulk soil (p = 0.0002, R^2^ = 0.51, Fig 3E, Table 1). The latter subsets did, however, show significantly different group dispersions (p < 0.05). The procedure used to identify OTUs that had significantly higher proportional representations in the N_0_P_0_ root data subset than in the other two (N_45_P_0_ root and N_0_P_0_ bulk soil) highlighted representatives of several taxa (Fig 4A).

**Fig 3.**
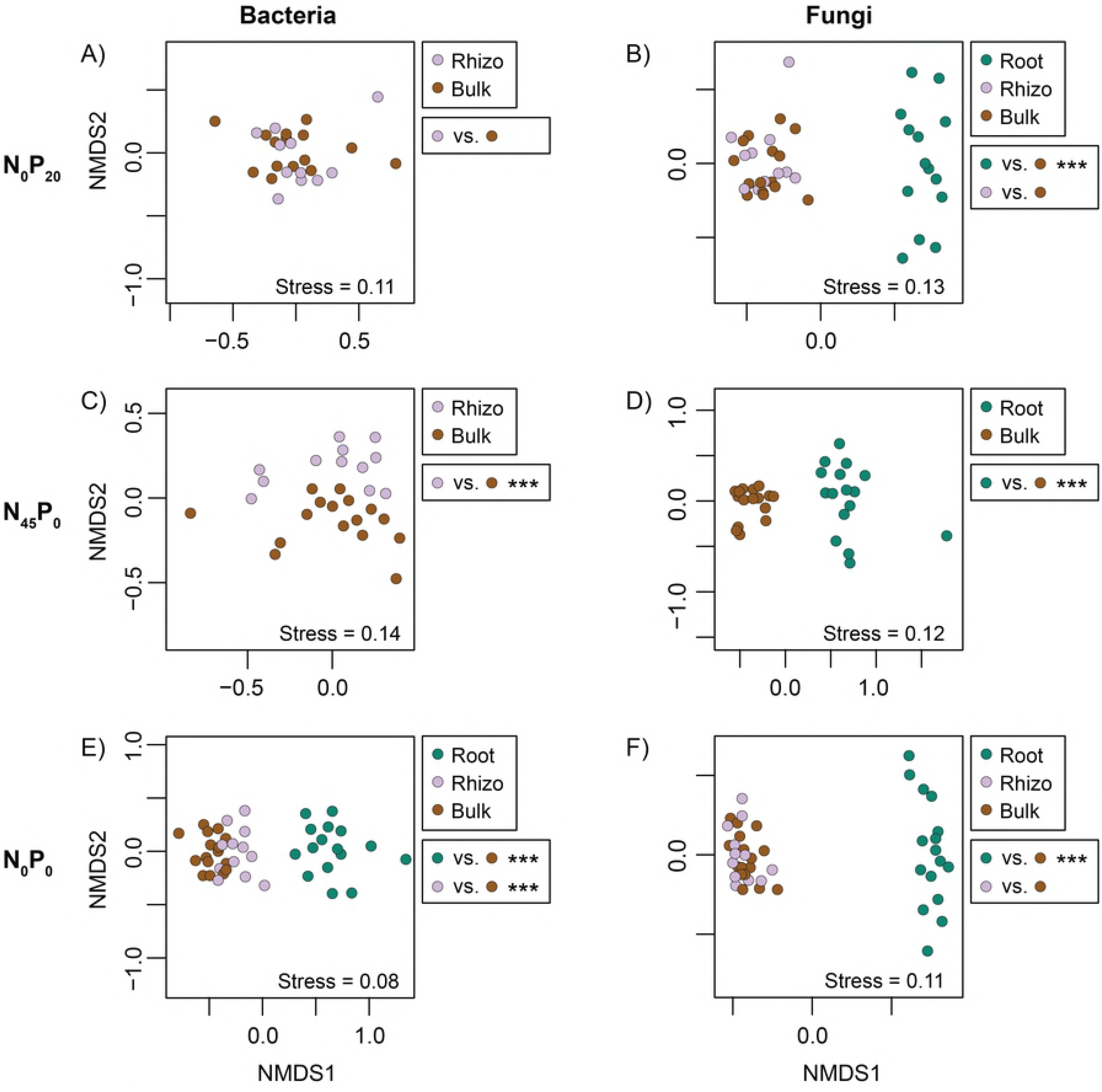
Microbial community comparisons - compartments. Non-metric multidimensional scaling (NMDS) ordinations representing the dissimilarity (Bray-Curtis) of microbial communities sampled from different soil/plant compartments. The significance of pairwise comparisons is reported next to each graph (* = p < 0.05; ** = p < 0.01, *** = p < 0.001).

**Table 1.** Summary of tests that allowed the identification of starvation- and *T. aestivum*-specific communities.

| Community | Compartment | Nutrient-sufficient soil | Nutrient-deficient soil | Treatment | Sufficient vs. starved (H1) | Starved vs. bulk soil deficient (H2) | Bulk soil starved vs. bulk soil deficient |
| --- | --- | --- | --- | --- | --- | --- | --- |
| Bacteria | Root | N45P0 | N0P0 | -N | *** | *** | *** |
| Bacteria | Rhizosphere | N45P0 | N0P0 | -N | *** | *** | *** |
| Fungi | Root | N45P20 | N0P20 | -N | ** | *** | *** |
| Fungi | Root | N45P0 | N0P0 | -N | *** | *** | *** |
| Bacteria | Rhizosphere | N45P20 | N45P0 | -P | * | *** |  |
| Fungi | Root | N45P20 | N45P0 | -P | ** | *** |  |

**Fig 4.**
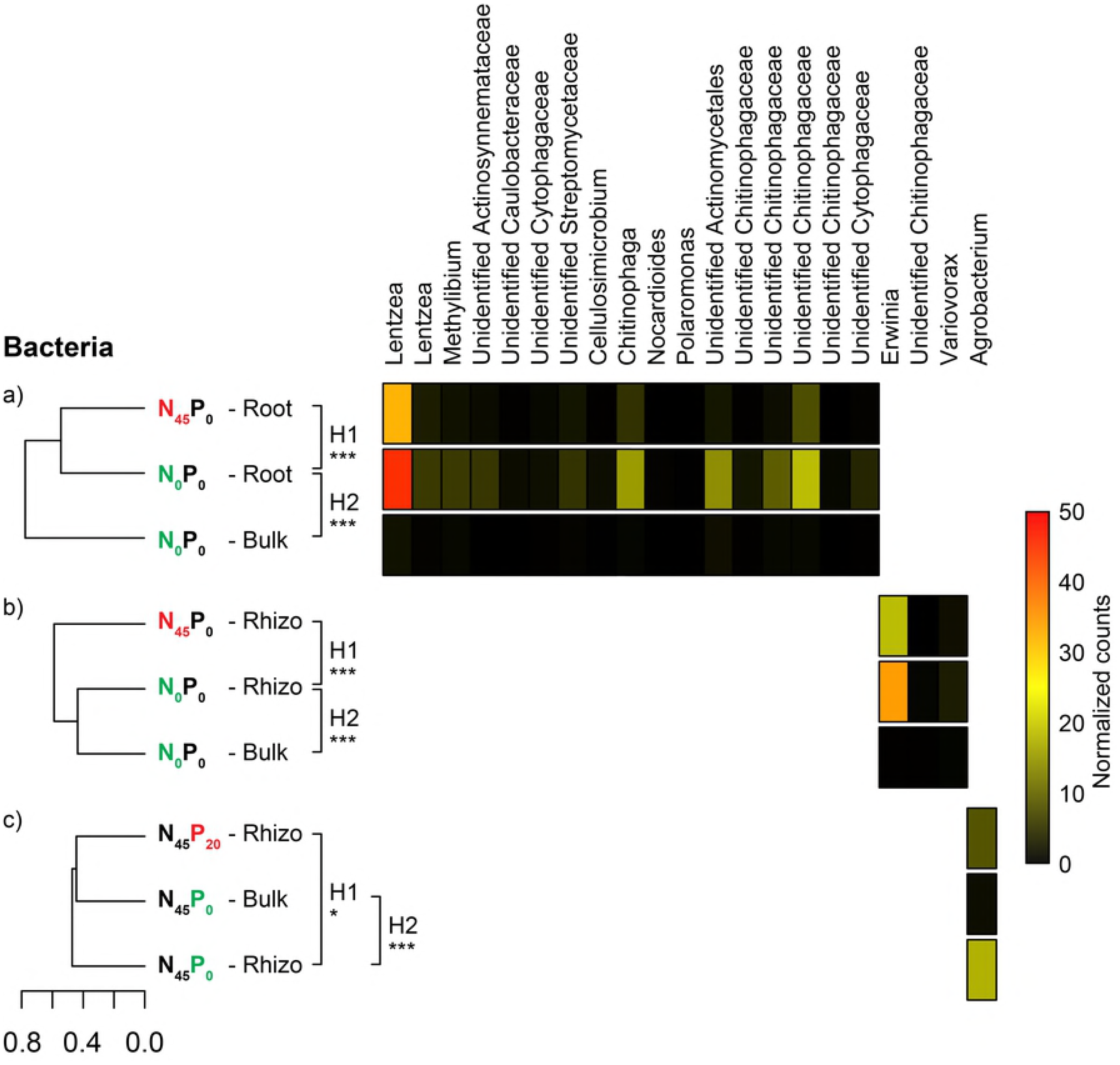
Bacterial OTU enrichments. Average dissimilarity (Bray-Curtis) of sets of compared bacterial communities, and relative abundance of OTUs that were significantly enriched in starvation- and *T. aestivum*-specific communities. (A) Data highlighting the singularity of N_0_P_0_ root communities, (B) data highlighting the singularity of N_0_P_0_ rhizosphere communities, (C) data highlighting the singularity of N_45_P_0_ rhizosphere communities. H1 = hypothesis 1: the nutrient-starved *T. aestivum* show root and rhizosphere microbial communities that are significantly different from those of the fertilized plants. H2 = hypothesis 2: starvation-specific *T. aestivum* communities are significantly different from those of the adjacent bulk soil. The significance of pairwise comparisons is reported next to each dendrogram (* = p < 0.05; ** = p < 0.01, *** = p < 0.001).

The bacterial rhizosphere communities of the four Rotation A plots showed significant differences of mean dissimilarity (p = 0.0001, R^2^ = 0. 27). The differences found in two pairs of N-starved/N-fertilized data subsets (N_45_P_20_/N_0_P_20_: p = 0.0004 and R^2^ = 0.18, N_45_P_0_/N_0_P_0_: p = 0.0004 and R^2^ = 0.27, Fig 2C) and one pair of P-starved/P-fertilized data subsets (N_45_P_20_/N_45_P_0_: p = 0.0176 and R^2^ = 0.12; Fig 2C) contributed to this signal. Only two of the latter three nutrient-starved subsets were also significantly different from their bulk soil counterparts (N_0_P_20_: p = 0.2649 and R^2^ = 0.07, N_45_P_0_: p = 0.0003 and R^2^ = 0.15, N_0_P_0_: p = 0.0003 and R^2^ = 0.17; Figs 3A, C, and E; Table 1). None of the above comparisons were performed between data subsets that showed significant differences of group dispersion. The procedure used to identify OTUs that had significantly higher proportional representations in the N_0_P_0_ rhizosphere subset than in the other two to which it was compared (N_45_P_0_ rhizosphere and N_0_P_0_ bulk soil) highlighted representatives of taxa *Erwinia, Chitinophagaceae*, and *Variovorax* (Fig 4B). The procedure used to identify OTUs that had significantly higher proportional representation in the N_45_P_0_ rhizosphere subset than in the other two to which it was compared (N_45_P_20_ rhizosphere and N_45_P_0_ bulk soil) highlighted a representative of genus *Agrobacterium* (Fig 4C).

Significant differences of mean community dissimilarity were found among the fungal root data of the four Rotation A plots (p = 0.0001, R^2^ = 0. 13). The differences found in two pairs of N-starved/N-fertilized data subsets (N_45_P_20_/N_0_P_20_: p = 0.0032 and R^2^ = 0.08, N_45_P_0_/N_0_P_0_: p = 0.0004 and R^2^ = 0.11, Fig 2B) and one pair of P-starved/P-fertilized data subsets (N_45_P_20_/N_45_P_0_: p = 0.0024 and R^2^ = 0.07; Fig 2B) contributed to this signal. All of the latter nutrient-starved subsets were also significantly different from their bulk soil counterparts (N_0_P_20_: p = 0.0003 and R^2^ = 0.28, N_45_P_0_: p = 0.0003 and R^2^ = 0.24, N0P0: p = 0.0003 and R^2^ = 0.22; Figs 3B, D, and F; Table 1). These comparisons were, however, performed on subsets that showed significantly different dispersions (p < 0.05). The procedure used to identify OTUs that had significantly higher proportional representations in the N_0_P_20_ root subset than in the other two to which they were compared (N_45_P_20_ root and N_0_P_20_ bulk soil) highlighted representatives of taxa *Lulworthia* and *Sordariomycetes* (Fig 5A). The procedure used to identify OTUs that had significantly higher proportional representation in the N_0_P_0_ root subset than in the other two to which they were compared N_45_P_0_ root and N_0_P_0_ bulk soil) highlighted a representative of taxa *Sordariomycetes, Apodus, Conocybe, Ascomycota, Crocicreas* (Fig 5B). The procedure used to identify OTUs that had significantly higher proportional representation in the N_45_P_0_ root subset than in the other two to which they were compared (N_45_P_20_ root and N_45_P_0_ bulk soil) highlighted a representative of genera *Parastagonospora* and *Phaeosphaeriopsis* (Fig 5C).

**Fig 5.**
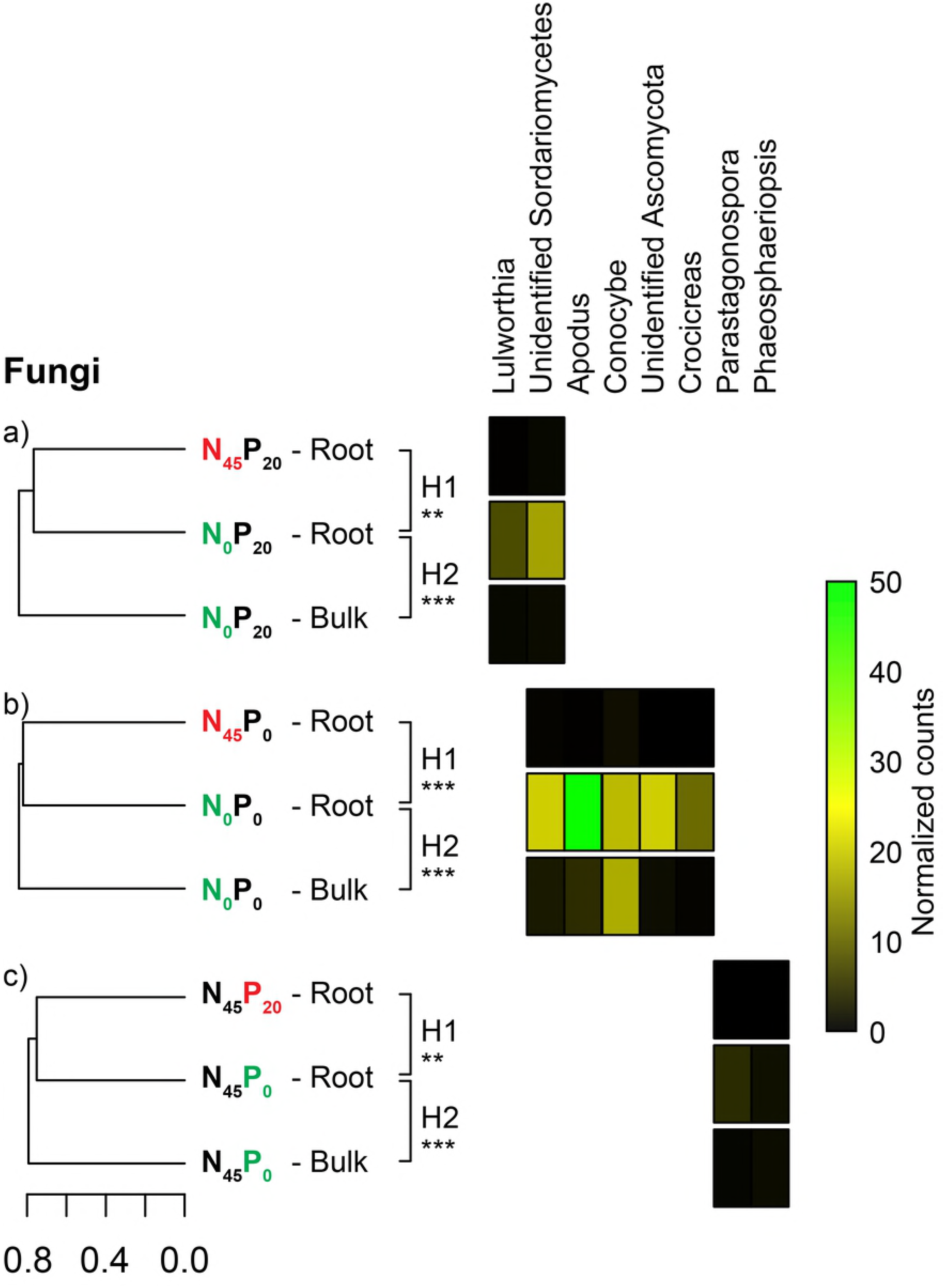
Fungal OTU enrichments. Average dissimilarity (Bray-Curtis) of sets of compared fungal communities, and relative abundance of OTUs that were significantly enriched in starvation- and *T. aestivum*-specific communities. (A) Data highlighting the singularity of N_0_P_20_ root communities, (B) data highlighting the singularity of N_0_P_0_ root communities, (C) data highlighting the singularity of N_45_P_0_ root communities. H1 = hypothesis 1: the nutrient-starved *T. aestivum* show root and rhizosphere microbial communities that are significantly different from those of the fertilized plants. H2 = hypothesis 2: starvation-specific *T. aestivum* communities are significantly different from those of the adjacent bulk soil. The significance of pairwise comparisons is reported next to each dendrogram (* = p < 0.05; ** = p < 0.01, *** = p < 0.001).

The fungal rhizosphere communities of the four Rotation A plots showed significant differences of mean dissimilarity (p = 0.0001, R^2^ = 0. 21). The difference found in two pairs of N-starved/N-fertilized data subsets (N_45_P_20_/N_0_P_20_: p = 0.0012 and R^2^ = 0.18, N_45_P_0_/N_0_P_0_: p = 0.0008 and R^2^ = 0.17, Fig 2D) contributed to this signal. However, neither of the nutrient-starved subsets were also significantly different from their bulk soil counterparts (N_0_P_20_: p = 0.2322 and R^2^ = 0.06, N_0_P_0_: p = 0.1728 and R^2^ = 0.06; Fig 3B, F; Table 1). None of the above comparisons were performed between data subsets that showed significant differences of group dispersion.

Significant differences of mean dissimilarity were, finally, also found among the bacterial and fungal bulk soil data of the four Rotation A plots (respectively p = 0.0001, R^2^ = 0.21 and p = 0.0001, R^2^ = 0.16). The differences found in pairs of N-starved/N-fertilized data subsets contributed to this signal in both cases (bacteria N_45_P_20_/N_0_P_20_: p = 0.0016 and R^2^ = 0.13, bacteria N_45_P_0_/N_0_P_0_: p = 0.0004 and R^2^ = 0.28, Fig 2E; fungi N_45_P_20_/N_0_P_20_: p = 0.0004 and R^2^ = 0.12, fungi N_45_P_0_/N_0_P_0_: p = 0. 0004 and R^2^ = 0.12, Fig 2F). None of the above comparisons were performed between data subsets that showed significant differences of group dispersion.

## Discussion

Our analyses identified four microbial communities that were specifically found in the roots or rhizospheres of N-starved *T. aestivum*. These communities were, of course, most probably shaped largely by factors that lie outside of plant-microbe partnerships. Their distinctiveness from the corresponding communities of N-fertilized plots, for one, was likely linked to the differences that existed between N45 and N0 bulk soil communities. Soil microbiome disparities are notoriously seen in agricultural fields that receive different N fertilization [33, 34, 63, 64]. They generally spread to plant microbiomes, and this seems to occur mostly through opportunistic microbial colonization [63, 64]. The four communities’ uniqueness could, however, also be partly tied to the activation of N-sharing plant-microbe partnerships in the N_0_ plots. Plant-associated communities that are molded by such interactions generally do, indeed, show N-dependent variations. Ikeda et al. [26, 27] reported on this phenomenon in soybeans. Others have similarly reported on correlations between the plant colonization efficiency of known N-providing plant partners and N soil content [22, 24]. This idea is also consistent with the additional differences noted between the four communities and their bulk soil counterparts. The latter pattern is indeed also seen in plant-associated communities that are molded by N-sharing partnerships. Zgadzaj et al. [36], for example, linked root-soil and rhizosphere-soil microbiome differences to the activation of microbial partner attraction and accommodation mechanisms in *Lotus japonicus*.

Many N-sharing plant-microbe partnerships have been identified outside of the *Fabaceae* family [42, 65, 66]. Others also seem to exist in plants like rice, *Arabidopsis thaliana*, and eucalyptus, which also show N-dependent microbiome variations [28, 30, 31, 43]. If they exist, *T. aestivum*’s N-sharing partnerships could be analogous. The microorganisms that were involved in the N0 plots were, however, likely not the most common N-providing plant partners. Indeed, none of the OTUs that were enriched in the four N-starvation and plant-specific communities belong to orders *Rhizobiales, Rhodospirillales, Cyanobacteriales*, or *Glomerales*. Several of the enriched OTUs do, however, belong to bacterial taxa that contain known diazotrophic plant partners (i.e. genus *Erwinia*, family *Enterobacteriaceae*, order *Burkholderiales*, class *Actinobacteria)* [42, 65]. Several others belong to taxa that were enriched in the microbiomes of N-deficient plants in other experiments (i.e. order *Burkholderiales*, families *Cytophagaceae, Chitinophagaceae, Caulobacteraceae*, and *Comamonadaceae)* [28, 30]. These OTUs may represent soil microbes that are specifically attracted towards *T. aestivum* when it experiences N-starvation, a behavior that is seen in known N-providing plant partners [46, 49].

The above observations do not constitute evidence for the existence of undiscovered N-sharing wheat-microbe partnerships, but they are coherent with it. Other such observations include links between the plant’s cultured root microbiome and soil N content [15], links between the plant’s metabolomic profile and soil N content [16, 20], the fact that it can shape its root microbiome through jasmonic acid production [17], the fact that its rhizosphere contains putative N-fixing microorganisms [18], and the fact that *T. aestivum* can directly receive N from microbes in experimental conditions [19]. Interestingly, several taxa that were represented on the list of OTUs enriched in the N-starvation and plant-specific communities contain known plant growth promoting microbes of other types. This is the case of bacterial genera *Methylibium, Polaromonas, Variovorax, Erwinia*, bacterial family *Chitinophagaceae*, which all contain known ACC deaminase producers [67–70], and bacterial genera *Cellulosimicrobium*, bacterial order *Actinomycetales*, which contain known antibiotic producers [71–73]. N starvation may thus also trigger non-N-sharing *T. aestivum-* microbe partnerships.

Our analyses also identified two microbial communities that were specifically found in the roots or rhizospheres of P-starved *T. aestivum*. These communities were also most probably shaped largely by factors that lie outside of plant-microbe partnerships. Their associated bulk soil communities were not, however, different from those associated with their P20 counterparts. In light of research that demonstrated correlations between plant microbiome composition and soil P content [32], correlations between the plant colonization efficiency of known P-providing plant partners and P soil content [23, 25, 29], and the root microbiome defining effects of microbial partner accommodation mechanisms in *A. thaliana* [44], these results suggest that the two communities may have been partly shaped by P-sharing plant-microbe partnerships. Wheat is a mycorrhizal plant [12–14], so the identification of a P-starvation- and *T. aestivum*-specific fungal root community was expected. The OTUs that were enriched in that community do not, however, belong to taxa known to contain fungal wheat partners (e.g. genera *Glomus, Sclerocystis, Acaulospora, Scutellospora)* or any other P-providing fungal plant partner (i.e. *Glomeromycetes, Agaricomycetes)* [41]. They rather belong to two taxa that contain notorious plant pathogens: genera *Phaeosphaeriopsis* and *Parastagonospora* [74, 75]. This observation is consistent with a P starvation-specific depression of *T. aestivum* immunity, a phenomenon that could be similar to that described in *A. thaliana* by Castrillo et al. [44]. The OTU that was enriched in the P-starvation- and *T. aestivum*-specific bacterial rhizosphere community does, however, belong to a genus known to contain phosphate-solubilizing plant growth promoters [76, 77].

## Conclusions

Six N/P starvation- and plant-specific microbial communities were identified. The OTUs that were enriched in these communities may represent microbes that are specifically attracted towards *T. aestivum* when it experiences N– or P-starvation, a behavior that is seen in known plant partners [46, 48, 49]. Many do, indeed, belong to taxa containing relevant N/P-providing plant partners. These results are consistent with the existence of undiscovered N/P-sharing wheat-microbe associations. Additional research will be needed to validate this interpretation. But the work presented here provides a way forward. The identification of potential *T. aestivum* partners gives us a target list for subsequent relationship-assessing studies, which is in line with modern microbiome research efforts that promote the identification of potentially beneficial microbes and their use in experimental system manipulations [78–80]. Wheat farming currently consumes approximately 20% of the worldwide production of inorganic N and P fertilizers, the latter experiments could thus pave the way for the development of valuable complementary yield-boosting tools.

## Acknowledgments

The authors acknowledge Julie Champagne (NRC Montreal), Miria Elias (NRC Montreal), Christine Maynard (NRC Montreal), and Danielle Ouellette (NRC Montreal) for the work they performed towards the production of 16S rRNA gene and ITS sequence data. The authors also thank Francis Gaudreault and Jessica Wasserscheid for their help with bioinformatics tools. We acknowledge Agriculture and Agri-Food Canada for access to samples obtained from the Historical Dryland Plots.

## Supporting information

**S1 Table. Sample summary.**

## References

1. Robertson GP, Vitousek PM. Nitrogen in agriculture: balancing the cost of an essential resource. Annual review of environment and resources. 2009 Nov 21;34:97–125.

2. Reay DS, Davidson EA, Smith KA, Smith P, Melillo JM, Dentener F, et al. Global agriculture and nitrous oxide emissions. Nature climate change. 2012 Jun;2(6):410.

3. Grassini P, Eskridge KM, Cassman KG. Distinguishing between yield advances and yield plateaus in historical crop production trends. Nature communications. 2013 Dec 17;4:2918.

4. Sharpley A. Managing agricultural phosphorus to minimize water quality impacts. Scientia Agricola. 2016 Feb;73(1):1–8.

5. Van Vuuren DP, Bouwman AF, Beusen AH. Phosphorus demand for the 1970-2100 period: a scenario analysis of resource depletion. Global environmental change. 2010 Aug 1;20(3):428–39.

6. Steffen W, Richardson K, Rockström J, Cornell SE, Fetzer I, Bennett EM, et al. Planetary boundaries: Guiding human development on a changing planet. Science. 2015 Feb 13;347(6223):1259855.

7. Fukuda M, Nagumo F, Nakamura S, Tobita S. Alternative fertilizer utilizing methods for sustaining low input agriculture. Soil Fertility 2012. InTech.

8. Calvo P, Nelson L, Kloepper JW. Agricultural uses of plant biostimulants. Plant and soil. 2014 Oct 1;383(1-2):3–41.

9. Chen X, Cui Z, Fan M, Vitousek P, Zhao M, Ma W, et al. Producing more grain with lower environmental costs. Nature. 2014 Oct;514(7523):486.

10. Han M, Okamoto M, Beatty PH, Rothstein SJ, Good AG. The genetics of nitrogen use efficiency in crop plants. Annual Review of Genetics. 2015 Nov 23;49:269–89.

11. Sala S, McLaren SJ, Notarnicola B, Saouter E, Sonesson U. In quest of reducing the environmental impacts of food production and consumption. Journal of cleaner production. 2017 Jan 1;140:387–98.

12. Cade-Menun BJ, Berch SM, Bomke AA. Seasonal colonization of winter wheat in South Coastal British Columbia by vesicular-arbuscular mycorrhizal fungi. Canadian journal of botany. 1991 Jan 1;69(1):78–86.

13. Li H, Smith SE, Holloway RE, Zhu Y, Smith FA. Arbuscular mycorrhizal fungi contribute to phosphorus uptake by wheat grown in a phosphorus-fixing soil even in the absence of positive growth responses. New Phytologist. 2006 Nov 1;172(3):536–43.

14. Duan J, Tian H, Drijber RA, Gao Y. Systemic and local regulation of phosphate and nitrogen transporter genes by arbuscular mycorrhizal fungi in roots of winter wheat (Triticum aestivum L.). Plant Physiology and Biochemistry. 2015 Nov 1;96:199–208.

15. Robinson RJ, Fraaije BA, Clark IM, Jackson RW, Hirsch PR, Mauchline TH. Endophytic bacterial community composition in wheat (Triticum aestivum) is determined by plant tissue type, developmental stage and soil nutrient availability. Plant and soil. 2016 Aug 1;405(1-2):381–96.

16. Heyneke E, Watanabe M, Erban A, Duan G, Buchner P, Walther D, et al. Characterization of the wheat leaf metabolome during grain filling and under varied N-supply. Frontiers in plant science. 2017 Nov 29;8:2048.

17. Liu H, Carvalhais LC, Schenk PM, Dennis PG. Effects of jasmonic acid signalling on the wheat microbiome differ between body sites. Scientific reports. 2017 Jan 30;7:41766.

18. Mahoney AK, Yin C, Hulbert SH. Community structure, species variation, and potential functions of rhizosphere-associated bacteria of different winter wheat (Triticum aestivum) cultivars. Frontiers in plant science. 2017 Feb 13;8:132.

19. Santos KF, Moure VR, Hauer V, Santos AS, Donatti L, Galvão CW, et al. Wheat colonization by an Azospirillum brasilense ammonium-excreting strain reveals upregulation of nitrogenase and superior plant growth promotion. Plant and Soil. 2017 Jun 1;415(1-2):245–55.

20. Zhang Y, Ma XM, Wang XC, Liu JH, Huang BY, Guo XY, et al. UPLC-QTOF analysis reveals metabolomic changes in the flag leaf of wheat (Triticum aestivum L.) under low-nitrogen stress. Plant Physiology and Biochemistry. 2017 Feb 1;111:30–8.

21. Heffer P, Prud’homme M. Global nitrogen fertilizer demand and supply: Trend, current level and outlook. International Nitrogen Initiative Conference. Melbourne, Australia 2016 Dec.

22. Carroll BJ, Mathews A. Nitrate inhibition of nodulation in legumes. Molecular biology of symbiotic nitrogen fixation. 1990:159–80.

23. Kabir Z, O’Halloran IP, Fyles JW, Hamel C. Seasonal changes of arbuscular mycorrhizal fungi as affected by tillage practices and fertilization: hyphal density and mycorrhizal root colonization. Plant and Soil. 1997 May 1;192(2):285–93.

24. Fuentes-Ramírez LE, Caballero-Mellado J, Sepúlveda J, Martínez-Romero E. Colonization of sugarcane by Acetobacter diazotrophicus is inhibited by high N-fertilization. FEMS Microbiology Ecology. 1999 Jun 1;29(2):117–28.

25. Treseder KK. A meta-analysis of mycorrhizal responses to nitrogen, phosphorus, and atmospheric CO2 in field studies. New Phytologist. 2004 Nov 1;164(2):347–55.

26. Ikeda S, Okubo T, Kaneko T, Inaba S, Maekawa T, Eda S, et al. Community shifts of soybean stem-associated bacteria responding to different nodulation phenotypes and N levels. The ISME journal. 2010 Mar;4(3):315.

27. Ikeda S, Anda M, Inaba S, Eda S, Sato S, Sasaki K, et al. Autoregulation of nodulation interferes with impacts of nitrogen fertilization levels on leaf-associated bacterial community in soybean. Applied and environmental microbiology. 2011 Jan 14.

28. Ikeda S, Sasaki K, Okubo T, Yamashita A, Terasawa K, Bao Z, et al. Low nitrogen fertilization adapts rice root microbiome to low nutrient environment by changing biogeochemical functions. Microbes and environments. 2014;29(1):50–9.

29. Liu Y, Shi G, Mao L, Cheng G, Jiang S, Ma X, et al. Direct and indirect influences of 8 yr of nitrogen and phosphorus fertilization on Glomeromycota in an alpine meadow ecosystem. New Phytologist. 2012 Apr;194(2):523–35.

30. Konishi N, Okubo T, Yamaya T, Hayakawa T, Minamisawa K. Nitrate supply-dependent shifts in communities of root-associated bacteria in Arabidopsis. Microbes and environments. 2017;32(4):314–23.

31. da Silva Fonseca E, Peixoto RS, Rosado AS, de Carvalho Balieiro F, Tiedje JM, da Costa Rachid CT. The microbiome of Eucalyptus roots under different management conditions and its potential for biological nitrogen fixation. Microbial ecology. 2018 Jan 1;75(1):183–91.

32. Pantigoso HA, Manter DK, Vivanco JM. Phosphorus addition shifts the microbial community in the rhizosphere of blueberry (Vaccinium corymbosum L.). Rhizosphere. 2018 Sep 1;7:1–7.

33. Hartmann M, Frey B, Mayer J, Mäder P, Widmer F. Distinct soil microbial diversity under long-term organic and conventional farming. The ISME journal. 2015 May;9(5):1177.

34. Sommermann L, Geistlinger J, Wibberg D, Deubel A, Zwanzig J, Babin D, et al. Fungal community profiles in agricultural soils of a long-term field trial under different tillage, fertilization and crop rotation conditions analyzed by high-throughput ITS-amplicon sequencing. PloS one. 2018 Apr 5;13(4):e0195345.

35. Oldroyd GE, Murray JD, Poole PS, Downie JA. The rules of engagement in the legume-rhizobial symbiosis. Annual review of genetics. 2011 Dec 15;45:119–44.

36. Zgadzaj R, Garrido-Oter R, Jensen DB, Koprivova A, Schulze-Lefert P, Radutoiu S. Root nodule symbiosis in Lotus japonicus drives the establishment of distinctive rhizosphere, root, and nodule bacterial communities. Proceedings of the National Academy of Sciences. 2016 Dec 6;113(49):E7996–8005.

37. Nishida H, Suzaki T. Nitrate-mediated control of root nodule symbiosis. Current opinion in plant biology. 2018 Aug 31;44:129–36.

38. Czarnecki O, Yang J, Weston DJ, Tuskan GA, Chen JG. A dual role of strigolactones in phosphate acquisition and utilization in plants. International journal of molecular sciences. 2013 Apr 9;14(4):7681–701.

39. Foo E, Ross JJ, Jones WT, Reid JB. Plant hormones in arbuscular mycorrhizal symbioses: an emerging role for gibberellins. Annals of botany. 2013 Mar 18;111(5):769–79.

40. Kapulnik Y, Koltai H. Fine-tuning by strigolactones of root response to low phosphate. Journal of integrative plant biology. 2016 Mar;58(3):203–12.

41. Johri AK, Oelmüller R, Dua M, Yadav V, Kumar M, Tuteja N, et al. Fungal association and utilization of phosphate by plants: success, limitations, and future prospects. Frontiers in microbiology. 2015 Oct 16;6:984.

42. Carvalho TL, Ballesteros HG, Thiebaut F, Ferreira PC, Hemerly AS. Nice to meet you: genetic, epigenetic and metabolic controls of plant perception of beneficial associative and endophytic diazotrophic bacteria in non-leguminous plants. Plant molecular biology. 2016 Apr 1;90(6):561–74.

43. Minamisawa K, Imaizumi-Anraku H, Bao Z, Shinoda R, Okubo T, Ikeda S. Are symbiotic methanotrophs key microbes for N acquisition in paddy rice root? Microbes and environments. 2016;31(1):4–10.

44. Castrillo G, Teixeira PJ, Paredes SH, Law TF, De Lorenzo L, Feltcher ME, et al. Root microbiota drive direct integration of phosphate stress and immunity. Nature. 2017 Mar;543(7646):513.

45. Yoneyama T, Terakado-Tonooka J, Minamisawa K. Exploration of bacterial N2-fixation systems in association with soil-grown sugarcane, sweet potato, and paddy rice: a review and synthesis. Soil Science and Plant Nutrition. 2017 Nov 2;63(6):578–90.

46. Sasse J, Martinoia E, Northen T. Feed your friends: do plant exudates shape the root microbiome? Trends in plant science. 2018 Oct, 23(1):25–41.

47. Janzen HH. The role of long-term sites in agroecological research: A case study. Canadian journal of soil science. 1995 Feb 1;75(1):123–33.

48. Nagahashi G, Douds DD. Isolated root caps, border cells, and mucilage from host roots stimulate hyphal branching of the arbuscular mycorrhizal fungus, Gigaspora gigantea. Mycological research. 2004 Sep;108(9):1079–88.

49. Vicré M, Santaella C, Blanchet S, Gateau A, Driouich A. Root border-like cells of Arabidopsis. Microscopical characterization and role in the interaction with rhizobacteria. Plant physiology. 2005 Jun 1;138(2):998–1008.

50. Siciliano SD, Germida JJ. Taxonomic diversity of bacteria associated with the roots of field-grown transgenic Brassica napus cv. Quest, compared to the non-transgenic B. napus cv. Excel and B. rapa cv. Parkland. FEMS Microbiology Ecology. 1999 Jul 1;29(3):263–72.

51. Huws SA, Edwards JE, Kim EJ, Scollan ND. Specificity and sensitivity of eubacterial primers utilized for molecular profiling of bacteria within complex microbial ecosystems. Journal of Microbiological Methods. 2007 Sep 1;70(3):565–9.

52. Martin KJ, Rygiewicz PT. Fungal-specific PCR primers developed for analysis of the ITS region of environmental DNA extracts. BMC microbiology. 2005 Dec;5(1):28.

53. Tremblay J, Singh K, Fern A, Kirton ES, He S, Woyke T, et al. Primer and platform effects on 16S rRNA tag sequencing. Frontiers in microbiology. 2015 Aug 4;6:771.

54. Magoč T, Salzberg SL. FLASH: fast length adjustment of short reads to improve genome assemblies. Bioinformatics. 2011 Sep 7;27(21):2957–63.

55. Edgar RC. Search and clustering orders of magnitude faster than BLAST. Bioinformatics. 2010 Aug 12;26(19):2460–1.

56. Wang Q, Garrity GM, Tiedje JM, Cole JR. Naive Bayesian classifier for rapid assignment of rRNA sequences into the new bacterial taxonomy. Applied and environmental microbiology. 2007 Aug 15;73(16):5261–7.

57. Robinson MD, McCarthy DJ, Smyth GK. edgeR: a Bioconductor package for differential expression analysis of digital gene expression data. Bioinformatics. 2010 Jan 1;26(1):139–40.

58. Marcon E, Hérault B. entropart: An R package to measure and partition diversity. Journal of Statistical Software. 2015;67(8).

59. Oksanen J, Kindt R, Legendre P, O’Hara B, Stevens MH, Oksanen MJ. The Vegan Package - Community Ecology Package. R package version 2.0-9.

60. Shaffer JP. Multiple hypothesis testing. Annual review of psychology. 1995 Feb;46(1):561–84.

61. Bretz F, Westfall P, Hothorn T. Multiple comparisons using R. Chapman and Hall/CRC; 2016 Apr 19.

62. Kolde R. pheatmap: Pretty heatmaps [Software].

63. Zhu S, Vivanco JM, Manter DK. Nitrogen fertilizer rate affects root exudation, the rhizosphere microbiome and nitrogen-use-efficiency of maize. Applied Soil Ecology. 2016 Nov 30;107:324–33.

64. Kavamura VN, Hayat R, Clark IM, Rossmann M, Mendes R, Hirsch PR, Mauchline TH. Inorganic nitrogen application affects both taxonomical and predicted functional structure of wheat rhizosphere bacterial communities. Frontiers in microbiology. 2018;9.

65. Franche C, Lindström K, Elmerich C. Nitrogen-fixing bacteria associated with leguminous and non-leguminous plants. Plant and soil. 2009 Aug 1;321(1-2):35–59.

66. Hodge A, Storer K. Arbuscular mycorrhiza and nitrogen: implications for individual plants through to ecosystems. Plant and soil. 2015 Jan 1;386(1-2):1–9.

67. Glick BR, Todorovic B, Czarny J, Cheng Z, Duan J, McConkey B. Promotion of plant growth by bacterial ACC deaminase. Critical Reviews in Plant Sciences. 2007 Oct 23;26(5-6):227–42.

68. Han JI, Choi HK, Lee SW, Orwin PM, Kim J, LaRoe SL, et al. Complete genome sequence of the metabolically versatile plant growth-promoting endophyte Variovorax paradoxus S110. Journal of bacteriology. 2011 Mar 1;193(5):1183–90.

69. Madhaiyan M, Poonguzhali S, Senthilkumar M, Pragatheswari D, Lee JS, Lee KC. Arachidicoccus rhizosphaerae gen. nov., sp. nov., a plant-growth-promoting bacterium in the family Chitinophagaceae isolated from rhizosphere soil. International journal of systematic and evolutionary microbiology. 2015 Feb 1;65(2):578–86.

70. Sagar A, Thomas G, Rai S, Mishra RK, Ramteke PW. Enhancement of Growth and Yield Parameters of Wheat Variety AAI-W6 by an Organic Farm Isolate of Plant Growth Promoting Erwinia Species (KP226572). International Journal of Agriculture, Environment and Biotechnology. 2018 Feb 1;11(1):159–71.

71. El-Tarabily KA, Sivasithamparam K. Non-streptomycete actinomycetes as biocontrol agents of soil-borne fungal plant pathogens and as plant growth promoters. Soil Biology and Biochemistry. 2006 Jul 1;38(7):1505–20.

72. Hasegawa S, Meguro A, Shimizu M, Nishimura T, Kunoh H. Endophytic actinomycetes and their interactions with host plants. Actinomycetologica. 2006 Dec 25;20(2):72–81.

73. Nabti E, Bensidhoum L, Tabli N, Dahel D, Weiss A, Rothballer M, et al. Growth stimulation of barley and biocontrol effect on plant pathogenic fungi by a Cellulosimicrobium sp. strain isolated from salt-affected rhizosphere soil in northwestern Algeria. European Journal of Soil Biology. 2014 Mar 1;61:20–6.

74. Golzar H, Wang C. First report of Phaeosphaeriopsis glaucopunctata as the cause of leaf spot and necrosis on Ruscus aculeatus in Australia. Australasian Plant Disease Notes. 2012 Dec 1;7(1):13–5.

75. Gao Y, Faris JD, Liu Z, Kim YM, Syme RA, Oliver RP, et al. Identification and characterization of the SnTox6-Snn6 interaction in the Parastagonospora nodorum-wheat pathosystem. Molecular Plant-Microbe Interactions. 2015 May;28(5):615–25.

76. Leyval C, Berthelin J. Interactions between Laccaria laccata, Agrobacterium radiobacter and beech roots: Influence on P, K, Mg, and Fe mobilization from minerals and plant growth. Plant and Soil. 1989 Jun 1;117(1):103–10.

77. Leyval CO, Surtiningsih TI, Berthelin JA. Mobilization of P and Cd from rock phosphates by rhizospheric microorganisms (phosphate-dissolving bacteria and ectomycorrhizal fungi). Phosphorus, Sulfur, and Silicon and the Related Elements. 1993 Apr 1;77(1-4):133–6.

78. Schlaeppi K, Bulgarelli D. The plant microbiome at work. Molecular Plant-Microbe Interactions. 2015 Mar;28(3):212–7.

79. Mitter B, Pfaffenbichler N, Sessitsch A. Plant–microbe partnerships in 2020. Microbial biotechnology. 2016 Sep;9(5):635–40.

80. Toju H, Peay KG, Yamamichi M, Narisawa K, Hiruma K, Naito K, Fukuda S, Ushio M, Nakaoka S, Onoda Y, Yoshida K. Core microbiomes for sustainable agroecosystems. Nat Plants. 2018 Apr 30;4:247–57.

